# Biochemical Characterization of RecBCD Enzyme from An Antarctic *Pseudomonas* Species and Identification of Its Cognate Chi (χ) Sequence

**DOI:** 10.1101/266460

**Authors:** Theetha L. Pavankumar, Anurag Kumar Sinha, Malay K. Ray

**Author notes:** These authors contributed equally to this work (Theetha L. Pavankumar, Anurag Kumar Sinha). Department of Biology, University of Copenhagen, Ole Maaløes Vej 5, Copenhagen, Denmark (Anurag Kumar Sinha). **Address for correspondence:** Theetha L Pavankumar, Department of Microbiology and Molecular Genetics, One Shields Ave, University of California, Davis, CA-95616, USA, Malay K. Ray, CSIR-Centre for Cellular and Molecular Biology, Uppal Road, Hyderabad 500 007, INDIA.

## Abstract

*Pseudomonas syringae* Lz4W RecBCD enzyme, RecBCD^Ps^, is a trimeric protein complex comprised of RecC, RecB, and RecD subunits. RecBCD enzyme is essential for *P. syringae* growth at low temperature, and it protects cells from low temperature induced replication arrest. In this study, we show that the RecBCD^Ps^ enzyme displays distinct biochemical behaviors. Unlike *E. coli* RecBCD enzyme, the RecD subunit is indispensable for RecBCD^Ps^ function. The RecD motor activity is essential for the Chi-like fragments production in *P. syringae*, highlighting a distinct role for *P. syringae* RecD subunit in DNA repair and recombination process. Further, the ssDNA-dependent endonuclease activity is notably absent in RecBCD^Ps^ enzyme. Here, we demonstrate that the RecBCD^Ps^ enzyme recognizes a unique octameric DNA sequence, 5′-GCTGGCGC-3′ (Chi^Ps^) that attenuates nuclease activity of the enzyme when it enters dsDNA from the 3′-end. We propose that the reduced translocation activities manifested by motor-defective mutants cause cold sensitivity in *P. syrinage*; emphasizing the importance of DNA processing and recombination functions in rescuing low temperature induced replication fork arrest.

**Abbreviations:**

ATP: Adenosine triphosphate
DSB: double-strand break
‘Chi: Crossover hotspot instigator
Ni-NTA: Nitrio tri-acetic acid
TLC: thin layer chromatography
MMC: mitomycin C
UV light: Ultra violet
ABM: Antarctic bacterial medium
LB: Luria-Bertani medium

## INTRODUCTION

The RecBCD enzyme-mediated homologous recombination is a DNA repair pathway that ensures genome integrity by faithful repair of broken DNA in *E. coli* and many Gram-negative bacteria. The heterotrimeric RecBCD enzyme complex, comprised of RecB, RecC, and RecD subunits, is essential for double-strand breaks (DSBs) repair via homologous recombination and protects host cells from foreign DNAs and invading phages (1-4). DSBs are generated in cells by various exogenous and endogenous factors including the running of replication forks into preexisting lesions (5). RecBCD enzyme is a highly processive helicase and nuclease, which unwinds and degrades DNA strands asymmetrically from a blunt or nearly blunt dsDNA (2). Initially, RecBCD enzyme degrades 3′-ended DNA preferentially over the 5′-end of the DNA until it encounters a regulatory DNA sequence called ‘Chi’ (<u>C</u>rossover <u>h</u>otspot *i*nstigator; 5′-GCTGGTGG-3′ in *E. coli*) (2, 6, 7). Chi (*χ*) recognition switches RecBCD enzyme’s polarity of DNA degradation. It attenuates 3′→5′ nuclease activity but upregulates 5′→3′ nuclease activity resulting in the production of 3′-ended ssDNA tail (8). The RecBCD enzyme loads RecA onto the 3′-terminal ssDNA (9) facilitating the homologous pairing to form Holliday junction. The RuvAB/C complex further resolve these recombination DNA intermediates to promote DNA repair process via homologous recombination (10). The behavior of RecBCD enzyme was also studied using single molecule techniques (11). Single molecule analyses of *E. coli.* RecBCD enzyme revealed that it translocates on DNA at a much higher rate, before Chi and pauses at Chi (12). The Chi recognition switches lead motor subunit of the RecBCD enzyme, from fast to slow motor (RecD to RecB), resulting in the reduction of translocation rate by one-half after Chi (13).

Previously, we have shown that the RecBCD enzyme of Antarctic *Pseudomonas syringae* Lz4W (RecBCD^Ps^) is essential for the growth at low temperature (14). Growth at low temperature induces frequent replication arrest causing fork reversal mediated DNA damage in *P. syringae* Lz4W (15). The RecBCD^Ps^ enzyme thus rescues cells from replication-arrest dependent DNA damage enabling *P. syringae* Lz4W to grow at low temperature (15). Unlike in *E. coli,* the RecD subunit of *P. syringae* Lz4W is an indispensable subunit of RecBCD complex (16). The genetic analysis of ATP binding and nuclease defective mutants of RecBCD^Ps^ enzyme indicated that the ATP driven motor activities of both RecD and RecB motor subunits are critical for growth at low temperature, whereas the nuclease activity of holoenzyme is dispensable (14).

In this study, we performed a biochemical analysis of wild-type and mutant RecBCD^Ps^ enzymes to understand their biochemical role in protecting *P. syringae* Lz4W cells from low temperature induced DNA damage. Here, we report that the ATP dependent DNA unwinding, not the DNA degradation activity of RecBCD enzyme is critical for growth at low temperature. Besides, inactivation of ATPase activity of RecD and RecB motor subunits has impaired the DNA unwinding activity of RecBCD enzyme leading to cold-sensitive phenotype *in vivo.* Furthermore, we have identified the *P. syringae* Chi sequence (5′-GCTGGCGC-3′), which attenuates the RecBCD^Ps^ nuclease activity *in vitro.*

## RESULTS

### Purification of RecBCD complex and its mutant variants from *P. syringae* Lz4W

The *P. syringae recCBD* genes were overexpressed on pGL10 derived plasmids in Δ*recCBD* (LCBD) strain to obtain RecB, RecC (N-terminal His-tagged) and, RecD proteins in an equimolar ratio (13). The wild-type and mutant RecBCD enzymes (RecB^K28Q^CD, RecBCD^K229Q^, and RecB^D1118A^CD) were purified by two-step purification protocol as shown in the flowchart (Fig. 1A) using Ni-NTA affinity column chromatography and size exclusion chromatography. The purified fractions of RecBCD and mutants contained all the three protein subunits (RecB, RecC, and RecD) in a stoichiometric ratio (Fig. 1B). Silver nitrate staining of gels further confirmed the purity of protein fractions. However, silver staining of SDS-PAGE separated proteins showed an additional protein band of ~60 kDa size (Fig. 1C). The Mass spectrometric analysis of this protein band indicated that it belongs to HSP-60 family of chaperonin, GroEL. The appearance of GroEL as a contaminant during purification of RecBCD protein is also evidenced in *E. coli (17).* Nonetheless, the presence of RecB, RecC, and RecD subunits was further confirmed by Western analysis using the polyclonal antibodies specific to these proteins (Fig. S1).

**FIGURE 1.**
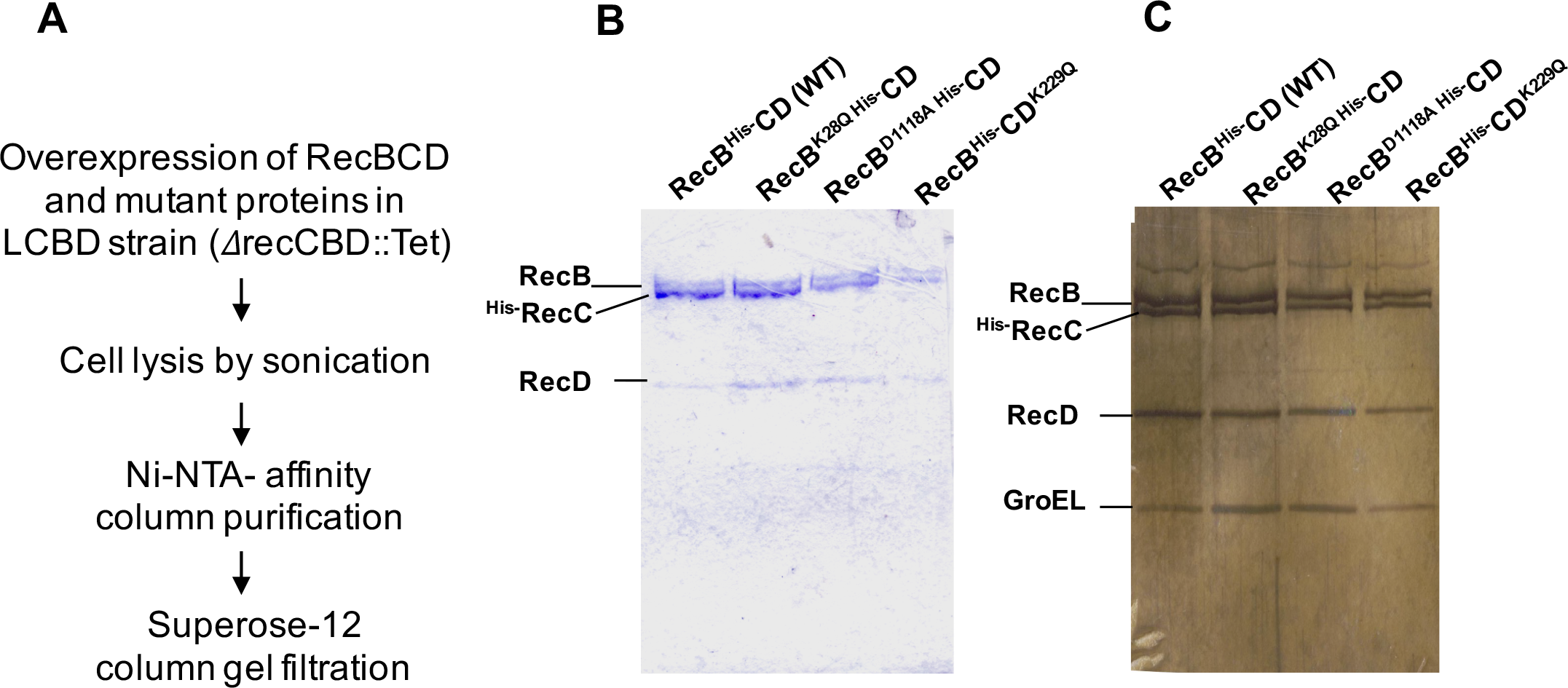
Purification of His-tagged RecBCD complex and mutant enzymes by Ni-NTA agarose column chromatography. **(A)** Steps involved in purification of wild-type RecBCD and mutant proteins. **(B)** SDS-PAGE analysis of purified RecBCD protein fractions of wild-type and mutants. Purified protein fractions were stained either with coomassie brilliant blue (B) or with silver nitrate (C). Three protein bands of expected size corresponding to RecB, RecC and RecD are visible on the gel. A low molecular protein (~60 kDa) observed on silver nitrate strained gel was identified as GroEL, a HSP-60 family chaperonin.

### ATP hydrolysis activity of the wild-type and the mutant RecBCD enzymes

ATP hydrolyzing activity of wild-type and mutant RecBCD enzymes was measured by thin layer chromatography (TLC) as described in Materials and methods. The wild-type RecBCD enzyme displayed DNA stimulated ATPase activity in the presence of linear pBR322 double-stranded DNA (dsDNA). At 22°C, RecBCD hydrolyzed ATP with the maximum velocity (V_max_) of 236.5 µmol ATP per µmol enzyme s^−1^ and a K_m_ of 57.8 µM for ATP (Fig. 2A and Table 1). The mutant RecB^D1118A^CD enzyme (nuclease defective mutant) showed ATPase activity similar to the wild-type enzyme. However, the ATP-binding defective mutant enzymes such as RecB^K28Q^CD (mutation in the consensus ATP binding site of RecB) and RecBCD^K229Q^ (mutation in the consensus ATP binding site of RecD) showed a 10-fold decrease in ATP hydrolyzing activities (Table 1).

**FIGURE 2.**
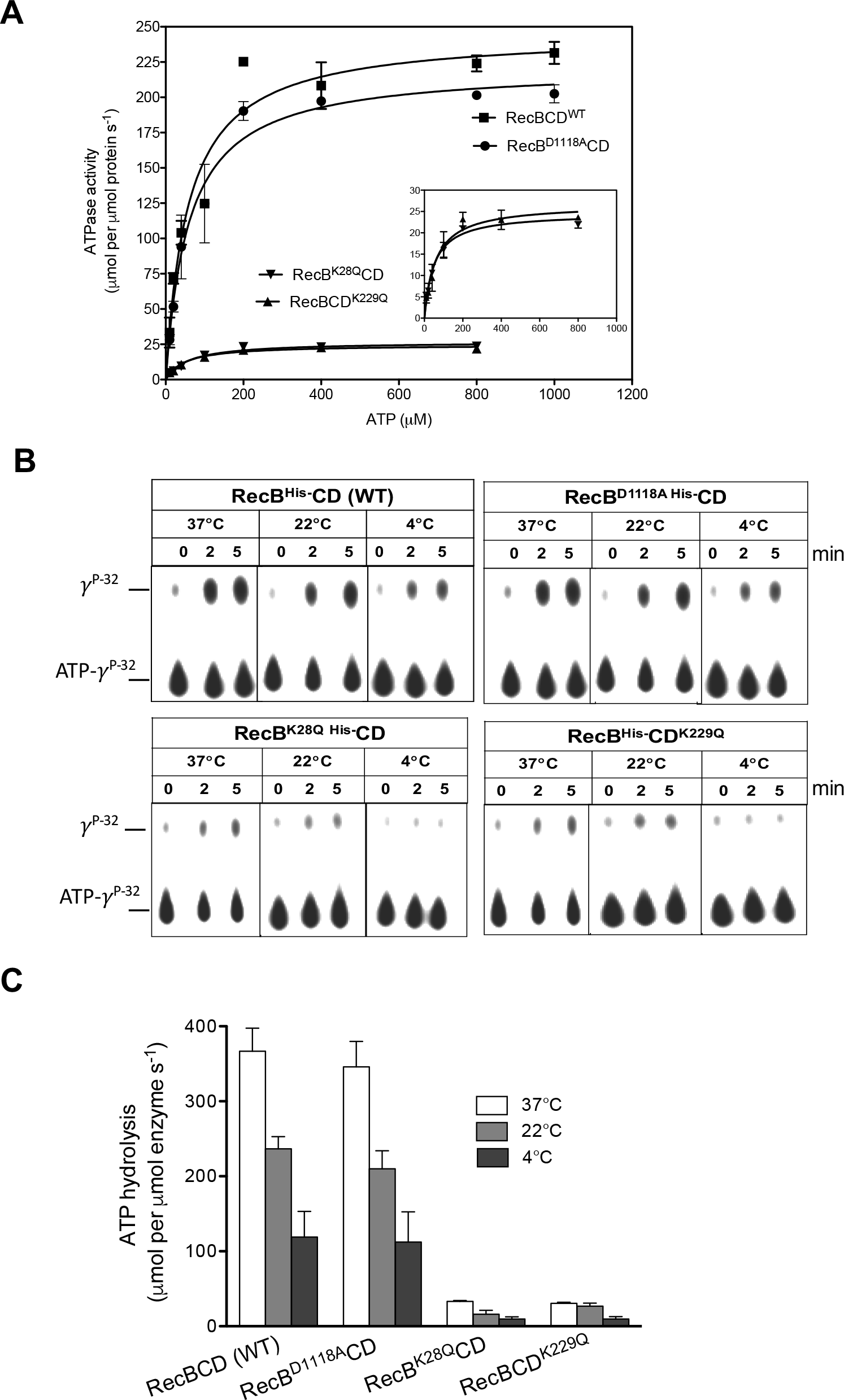
ATP hydrolysis of wild-type and mutant RecBCD enzymes at different temperatures. The assays were carried out by TLC method as described in Methods. **(A)** A graph showing the concentration dependent ATP hydrolysis by RecBCD (WT) and mutant enzymes at 22°C. The inset in B shows a blow-up of the same data of mutant enzymes using an expanded *γ*-scale. **(B)** Representative of TLC plates showing the ATP hydrolysis by wild-type and mutant RecBCD enzymes at 37°, 22° and 4°C. **(C)** ATPase activities at 37°, 22° and 4°C are shown in histogram with error bars. The ATP hydrolysis data presented in the histogram represents the results obtained from three independent experiments.

**TABLE 1.**
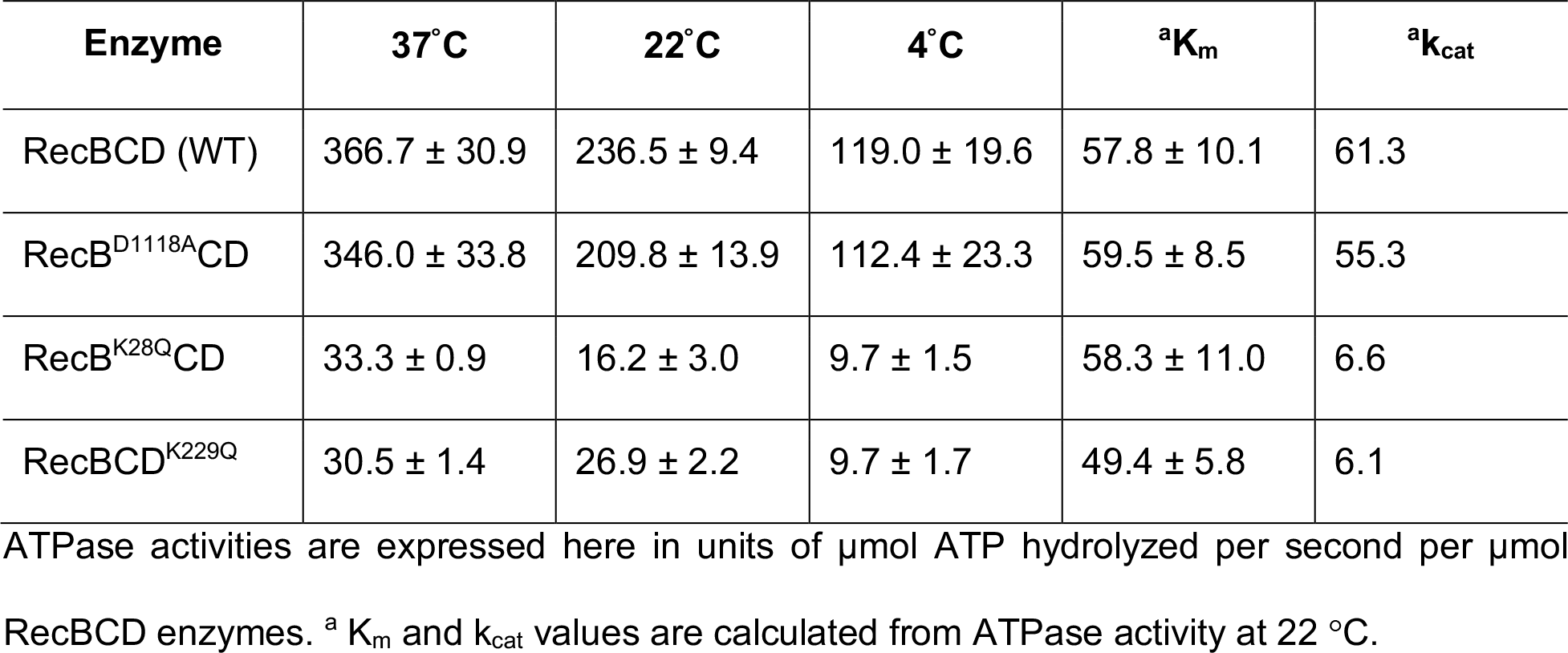
ATPase activity of wild type and mutant RecBCD enzymes

ATP hydrolyzing activities of RecBCD enzymes were measured at three different temperatures (37, 22 and 4°C) (Fig. 2B). Interestingly, The RecBCD enzyme showed the highest ATPase activity at 37°C compared to 22°C and 4°C. At 22°C, the optimum temperature for *P. syringae* Lz4W growth, the wild-type RecBCD and the nuclease-deficient RecB^D1118A^CD enzymes displayed ~40% lower ATPase activity compared to 37°C. At 4°C, the activities were further reduced to ~50% of their respective ATPase activity observed at 22°C. The ATP hydrolyzing defective mutants showed ~10-fold decrease in ATP hydrolyzing activity compared to wild-type RecBCD enzyme, at all the temperatures tested (Table 1 and Fig. 2C). It indicates that the mutations were previously shown to prevent *P. syringae* growth at low temperature severely curtailed ATP hydrolyzing activity.

### DNA unwinding and degradation activities of the RecBCD enzyme at different concentration of Mg^++^ and ATP

The DNA unwinding and degradation activity of *E. coli* RecBCD enzyme (RecBCD^Ec^) are free [Mg^2+^] ion dependent. An increase in free [Mg^2+^] increases nucleolytic cleavage by RecBCD enzyme (18). To understand the DNA unwinding and degradation behavior of *P. syringae* RecBCD enzyme (RecBCD^Ps^), we performed experiments at various concentrations of magnesium and ATP as described in Materials and methods. In the first set of reactions, the molar concentration of magnesium was kept constant (2 mM) and the ATP concentration was varied (0-10 mM); and in the second set of reactions, the ATP concentration was kept constant (2 mM) and magnesium concentration was varied (0-10 mM). The results indicate that the DNA unwinding and degradation properties of RecBCD^Ps^ enzyme are also modulated by the ratio of Mg^2+^ and ATP. When the molar concentration of Mg^2+^ is lesser than ATP, RecBCD^Ps^ enzyme unwinds the dsDNA substrate, and the degradation of DNA is not much pronounced (Fig. 3A, lane 4-6). However, when the molar concentration of Mg^++^ exceeds the ATP concentration, RecBCD^Ps^ degrades unwound DNA more vigorously (Fig. 3B, lane 4-10). The ratio of [Mg^2+^]: [ATP] thus affects the kinetics of DNA unwinding and degradation properties of RecBCD enzyme. We also observed that an increase in ATP concentration more than three folds over Mg^++^ inhibits the DNA unwinding activity of RecBCD^Ps^ enzyme (Fig. 3A, lane 7-10). The inhibition of DNA unwinding activity is possibly due to sequestration of a Mg^2+^ ion by ATP leading to the depletion of free magnesium in the reaction. Subsequently, the DNA unwinding and degradation experiments were performed under specified reaction conditions. The limiting magnesium reaction condition (5 mM ATP, 2 mM Mg^++^) was chosen to observe the DNA unwinding activity and, the excess magnesium condition (2 mM ATP, 6 mM Mg^++^) to observe the DNA unwinding and degradation activities of RecBCD enzyme. Interestingly, under excess magnesium conditions, we observed three shorter discrete DNA fragments (Fig. 3B, Lane 5-10). These DNA fragments production by RecBCD enzyme is possibly due to Chi-like sequence on a DNA substrate and is discussed later in the results section.

**FIGURE 3.**
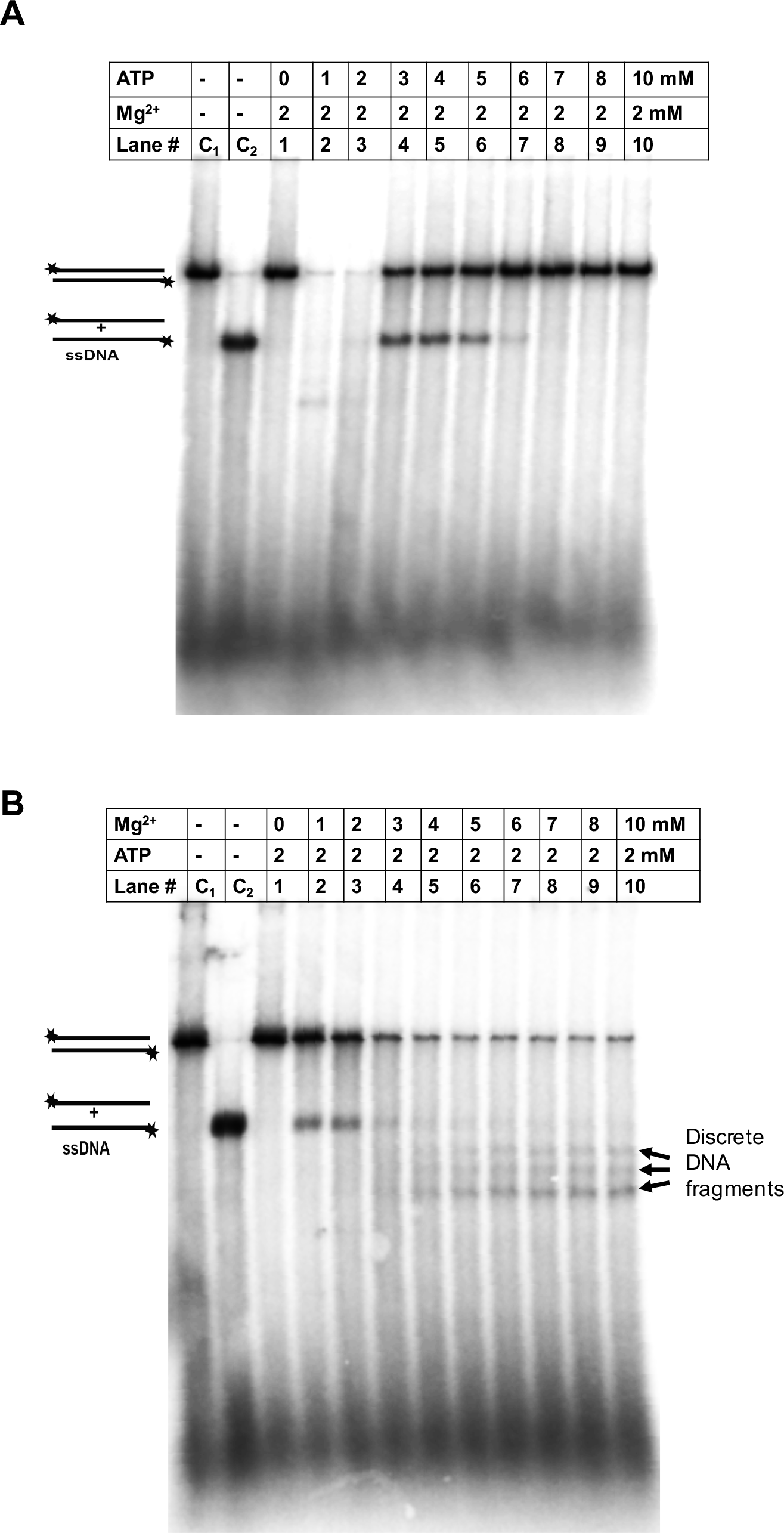
Effects of magnesium and ATP on the unwinding and degradation by wild-type RecBCD enzyme of *P. syringae.* The DNA unwinding and degradation assays were performed with [5′-^32^P] labeled *Nde*I linearized pBR322 plasmid DNA in the presence of different concentrations of magnesium and ATP, as described in Methods; **(A)** the reaction mixture contained fixed amount of Mg-acetate (2 mM) and the amount ATP was varied (0 to 10 mM) as indicated. **(B)** The reaction mixture contained the fixed amount of ATP (2 mM) and concentration of Mg-acetate was varied (0 to 10 mM) as indicated. The reactions were performed for 5 min and analyzed on a 1% agarose gel containing 1X TBE buffer at a 25-Volts for hrs. Agarose gels were dried, exposed to phosphor imaging plates and quantified using Phosphor Imager (Fuji-3000). The data were analyzed using image gauge software. The lanes C_1_ and C_2_ contain [5′-^32^P] labeled dsDNA and the heat denatured 5′-^32^P labeled dsDNA respectively as a control. A [5′-^32^P] labeled discrete DNA bands of smaller than full-length ssDNA of pBR322 were also noticed in the lanes 5-10 of panel B.

Calcium has been shown to inhibit the nuclease activity of *E. coli* RecBCD enzyme (19). We also studied the effects of calcium on *P. syringae* RecBCD by increasing the Ca^++^ ion concentration in a reaction mixture that contained fixed amounts of ATP and Mg^2+^ (2 and 6 mM respectively) (Supplementary Fig. S2). We found that similar to RecBCD^Ec^, the high concentration of calcium inhibits both helicase and nuclease activity of RecBCD^Ps^ enzyme.

### Temperature-dependent DNA unwinding and nuclease activity of *P. syringae* RecBCD enzyme

ATPase RecBCD mutants affect *P. syringae* Lz4W growth in a temperature-dependent manner. Hence, we sought to assess the effects of temperature on the DNA unwinding and degradation properties of RecBCD^Ps^ enzyme. We measured the enzyme activities at 22°C, and 4°C, using NdeI linearized [5′-^32^P] labeled pBR322 plasmid as a substrate. Under the limiting magnesium reaction condition (5 mM ATP, 2 mM Mg^2+^), the DNA unwinding activity (i.e., production of full-length ssDNA) of RecBCD^Ps^ enzyme was detected only at 22°C and not at 4°C (Fig. 4A). However, under excess magnesium reaction condition (2 mM ATP, 6 mM Mg^++^) the DNA degradation activity of RecBCD^Ps^ enzyme was observed at both 22°C and 4°C (Fig. 4B).

**FIGURE 4.**
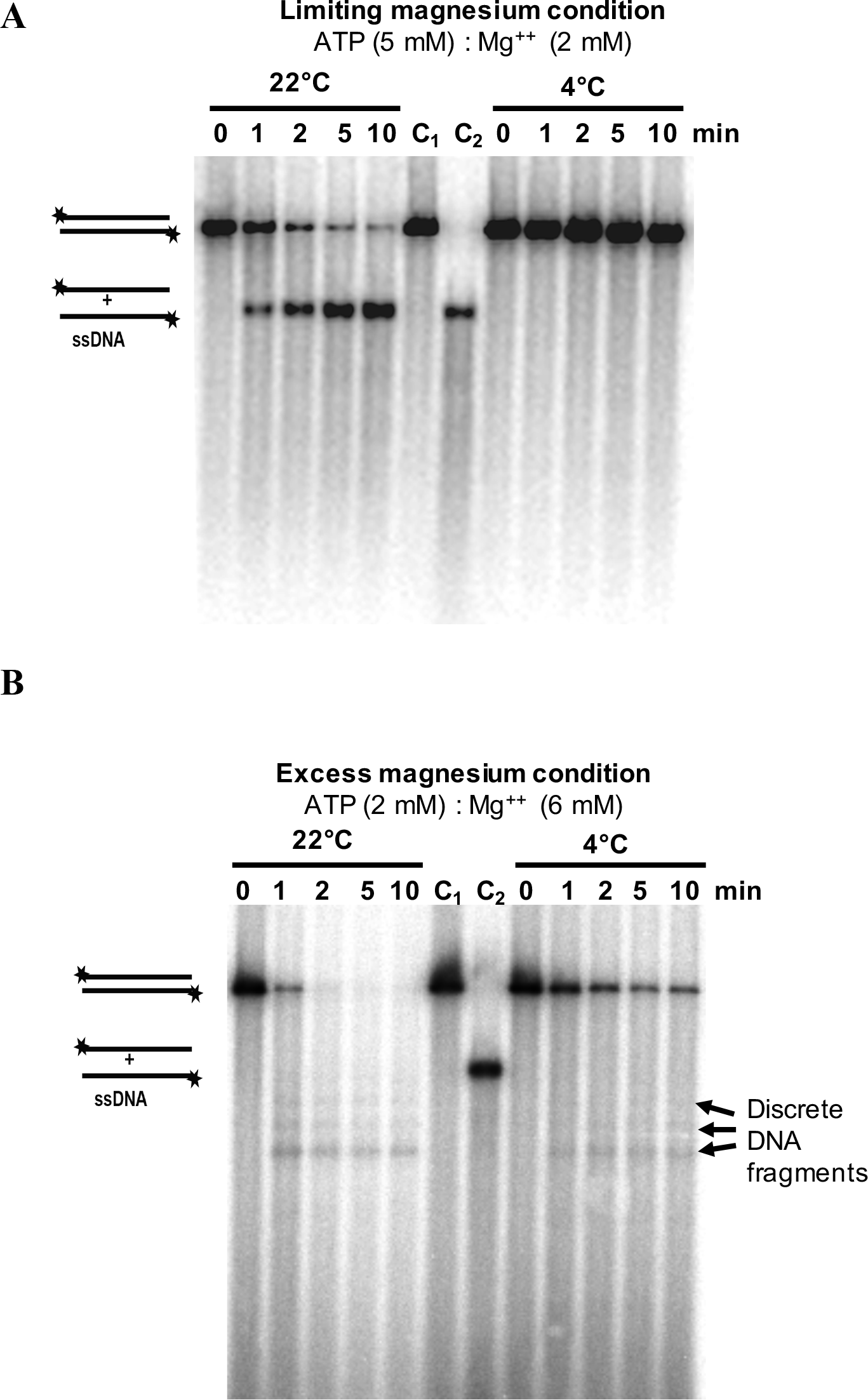
DNA unwinding and degradation of *NdeI* digested linear dsDNA of pBR322 by RecBCD^Ps^ (WT) enzyme at 22° and 4°C. The DNA unwinding and degradation at 22° and 4°C were carried out as described in Methods. **(A)** The DNA unwinding reactions contained 5 mM ATP and 2 mM Mg^++^ (limiting magnesium condition) **(B)** The DNA degradation reactions contained 2 mM ATP and 6 mM Mg^++^ (excess magnesium condition). The reactions were initiated by adding ATP, and stopped at the indicated times by adding stop-buffer. The lanes C_1_ and C_2_ contain [5′-^32^P] labeled *Nde*I linearised double-stranded and the heat-denatured ssDNA of pBR322 as control. The discrete ssDNA fragments of pBR322 produced by the nuclease activity of RecBCD enzyme are also indicated.

We calculated the rate of DNA unwinding by measuring the decreased band intensities of 5′-[*γ*-^32^P]-labeled dsDNA substrates at different temperatures. The wild-type RecBCD^Ps^ unwound the DNA at the rate of 35.1 ± 1.6 bp/sec at 22°C when ATP and Mg^++^ ratio was 5:2 (limiting magnesium condition) (Table 2). However, the enzyme could unwind the DNA even much faster, when the ATP and Mg^++^ ratio was changed to 2:6 (excess magnesium condition). Under the latter condition, wild-type RecBCD^Ps^ enzyme unwound (also degraded) the DNA at the rate of 101.5 ± 3.3 bp/sec (Table 2). Notably, under the limiting magnesium reaction condition at low temperature (4°C), the RecBCD enzyme failed to produce detectable unwound DNA products (Fig. 4A). However, under excess magnesium conditions, RecBCD enzyme could unwind and degrade the DNA at the rate of 55.8 ± 6.9 bp/s (Table 2, Fig. 4B), which is about 54% of the activity observed at 22°C. The apparent unwinding rates obtained under excess magnesium conditions, based on the disappearance of ^32^P-end-labeled DNA substrate could be an over-estimation. It is possible that some enzyme molecules could just nucleolytically remove the end-labeled nucleotide, but couldn’t fully unwind the dsDNA substrate.

**TABLE 2.**
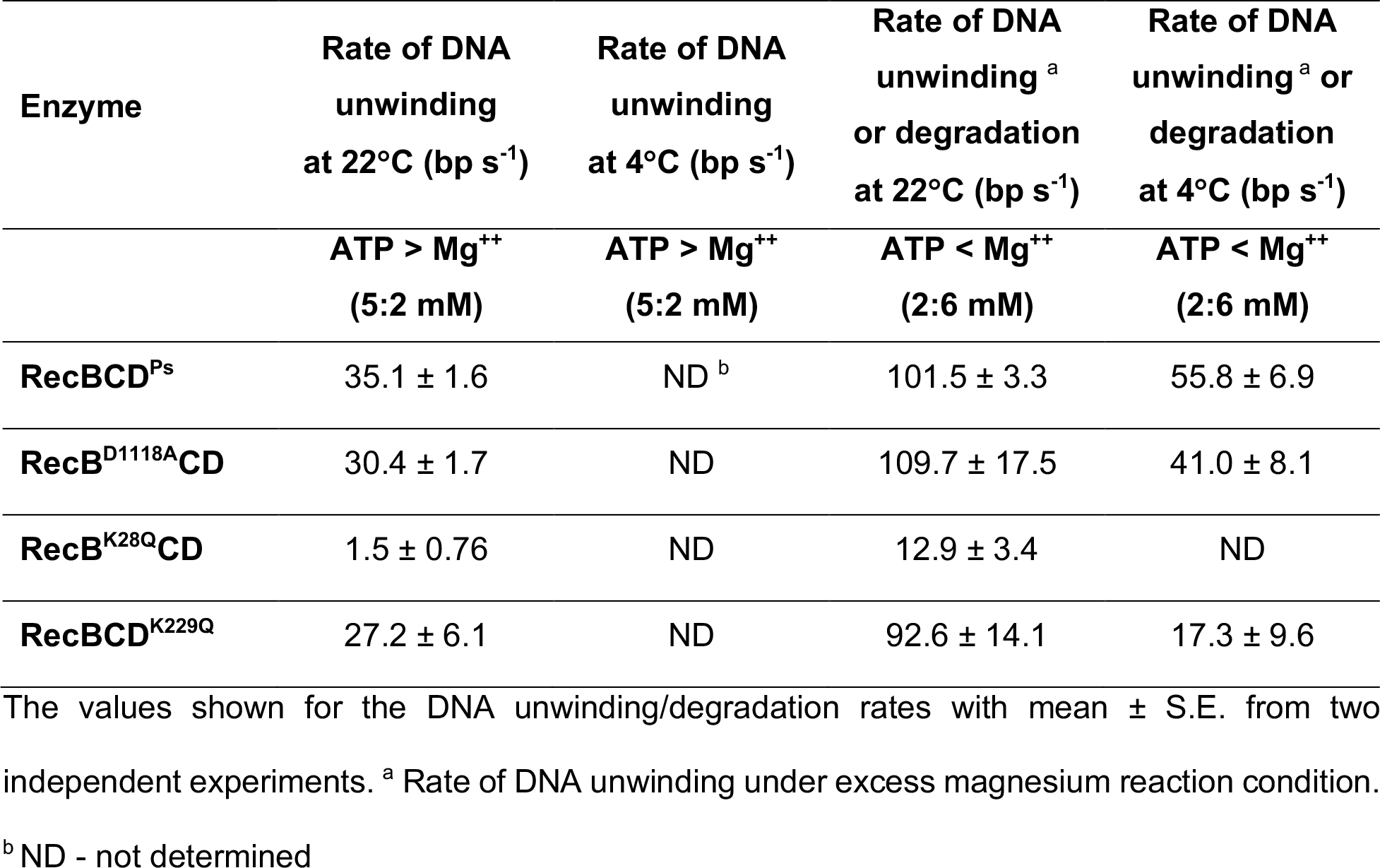
DNA unwinding and degradation activities of the wild type and mutant RecBCD enzymes at and 4°C

### DNA unwinding and nuclease activities of mutant RecB^K28Q^CD, RecBCD^K229Q^, and RecB^D1118A^CD enzymes

We have shown that *P. syringae* cells carrying *recB^K28Q^CD* or *recBCD^K229Q^* mutants are sensitive to cold temperature, UV irradiation and Mitomycin C (MMC) (14). These two mutant enzymes also display very weak ATPase activity. To understand the impact of these mutations on the DNA unwinding and degradation properties of RecBCD complex, we analyzed RecB^K28Q^CD and RecBCD^K229Q^ enzymes *in vitro* at 22°C and 4°C. At 22°C, under limiting and excess magnesium conditions, RecB^K28Q^CD with the ATPase-defective RecB subunit displayed weak helicase activity (Fig. 5A). The rate of DNA unwinding was 1.5 ± 0.7 bp/sec and 12.9 ± 3.4 bp/sec at 22°C, under limiting and excess magnesium conditions respectively (Fig.5A and Table 2). At 4°C, on the other hand, RecB^K28Q^CD showed no detectable DNA unwinding activity under the excess magnesium condition (Fig. 5A and Table 2). These results suggest that RecB^K28Q^CD is a poor helicase-nuclease enzyme.

**FIGURE 5.**
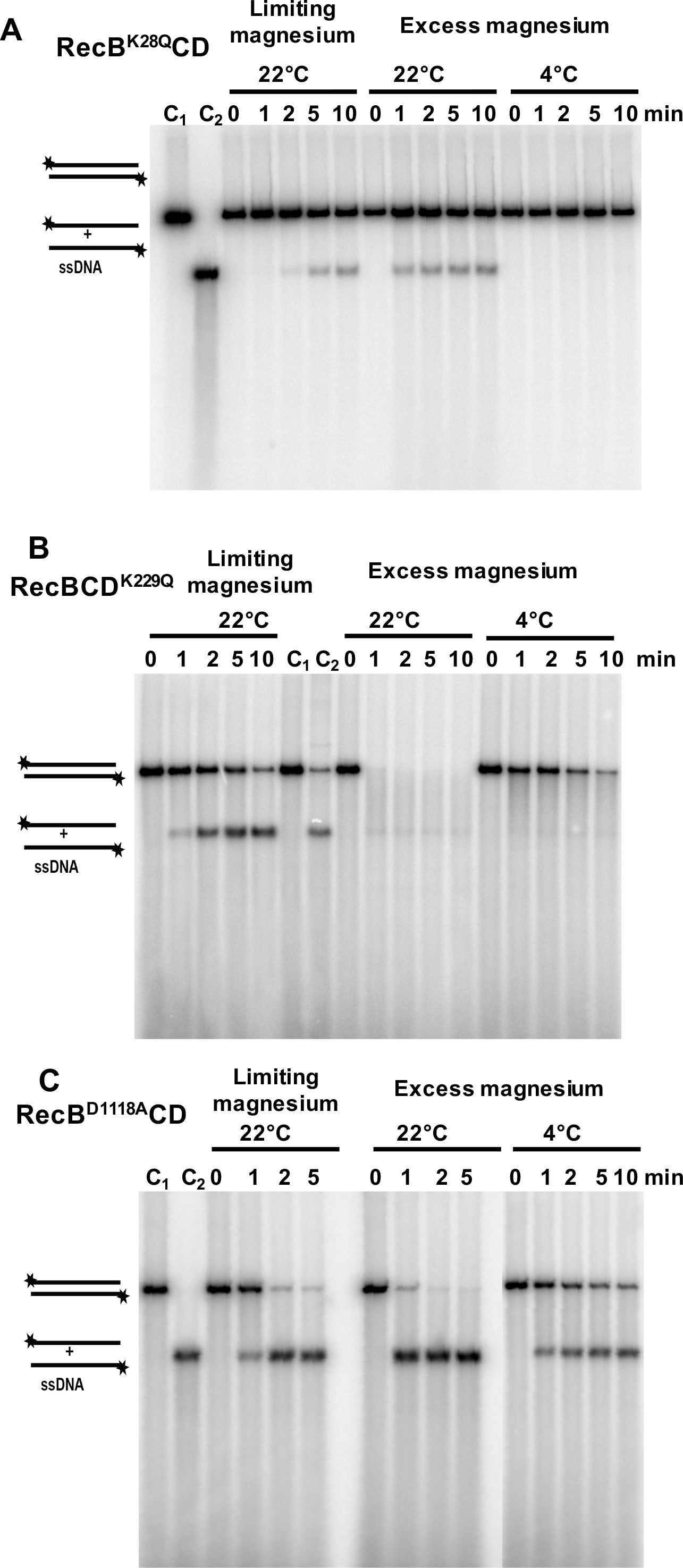
DNA unwinding and degradation of *NdeI* linearized dsDNA of pBR322 by mutant RecBCD^Ps^ enzymes at 22° and 4°C. (A) DNA unwinding and degradation by RecB^K28Q^CD enzyme. The DNA unwinding and degradation at 22° and 4°C were carried out as described in Methods. Note that this enzyme shows DNA unwinding (ssDNA production), but no detectable DNA degradation at 22°C, and none were detectable at 4°C. **(B) DNA unwinding and degradation by RecBCD^K229Q^ enzyme.** The RecBCD^K229Q^ enzyme has apparently retained both the DNA unwinding and degradation properties. But, interestingly, the discrete DNA bands are absent **(C) DNA unwinding by nuclease-deficient RecB^D1118A^CD enzyme under limiting and excess magnesium conditions at and 4°C.** RecB^D1118A^CD enzyme is unable to degrade DNA at both 22 and 4°C and notably, the DNA unwinding seems to be faster in excess-magnesium reaction condition (2 mM ATP:6 mM Mg^++^) compared to limiting magnesium reaction condition (5 mM ATP:2 mM Mg^++^). Each reaction mixtures contained 0.5 nM of enzyme and 10 µM (nucleotides) linear [5′-^32^P] labeled pBR322 dsDNA. The lanes C_1_ and C_2_ contain [5′-^32^P] labeled *NdeI* linearized double-stranded and the heat-denatured ssDNA of pBR322 respectively.

Compared to the RecB^K28Q^CD enzyme, RecBCD^K229Q^ enzyme with the ATPase-defective motor RecD subunit displayed higher DNA unwinding and nuclease activities. At 22°C, the RecBCD^K229Q^ enzyme with the ATPase-defective RecD subunit displayed substantial DNA unwinding activity, 27.2 ± 6.1 bp/sec and 92.6 ± 14.1 bp/sec, under limiting and excess magnesium conditions respectively (Fig. 5B and Table 2). However, at 4°C under excess magnesium conditions, the mutant enzyme unwound the DNA at the rate of 17.3 ± 9.6 bp/sec (Fig 5B and Table 2). The combined DNA unwinding and degradation activity of the RecBCD^K229Q^ enzyme (under excess magnesium) were ~90% and ~30% of the wild-type RecBCD^Ps^ activity at 22°C and 4°C respectively. Interestingly, the mutant RecBCD^K229Q^ enzyme failed to produce discrete DNA fragments (Fig. 5B).

We also tested DNA unwinding and degradation by the nuclease defective RecB^D1118A^CD enzyme. (Fig. 5C). The mutation in the nuclease catalytic site of RecB subunit (RecB^D1118A^CD) led to a complete loss of *in vivo* nuclease activity in RecBCD^Ps^ complex, without affecting the recombination proficiency and cold adaptation of bacteria (14, 15). At 22°C, under limiting magnesium reaction condition, RecB^D1118A^CD enzyme produced single-stranded pBR322 DNA at the rate 30.4 ± 1.7 bp/sec, which is 85% of the rate observed with the wild-type enzyme. Under excess magnesium condition, the rate of DNA unwinding increased to 109.7 ± 17.5 bp/sec, which is similar to wild-type. At 4°C, this enzyme could unwind the pBR322 DNA at the rate of 41.0 ± 8.1 bp/sec, which is about 75% of wild-type enzyme (Table 2). More importantly, under both limiting and excess magnesium conditions, this enzyme as expected, produced only the full-length ssDNA of pBR322 and did not degrade the DNA (Fig. 5C). From these data, it is clear that RecB^D1118A^CD enzyme is nuclease deficient *in vitro,* but its DNA unwinding activity is comparable to wild-type enzyme at both 22 and 4°C (Fig. 5C and Table 2).

### *P. syringae* RecBCD enzyme does not have endonuclease activity

The ATP-dependent endonuclease activity of wild-type RecBCD enzyme of *P. syringae* was examined using a circular M13 ssDNA as substrate as previously described for RecBCD^Ec^ (20-22). We tested the endonuclease activity under three different conditions of ATP and magnesium concentrations as described in Supplementary Fig S4. Under all the three conditions, wild-type RecBCD and mutant proteins failed to degrade M13 phage ssDNA (Fig S3). It indicates that unlike RecBCD^Ec^, the RecBCD^Ps^ does not exhibit endonucleolytic activity under the conditions tested.

### A specific DNA sequence on pBR322 plasmid modulates nuclease activity of *P. syringae* RecBCD

We noticed that, under the excess magnesium reaction conditions, RecBCD^Ps^ produced a discrete DNA fragments shorter than the full-length unwound DNA substrate (Fig 3B, Lanes 5-10), and the shortest DNA fragment showed a higher intensity than the other ones (Fig 3B). This interesting observation led us to hypothesize that the *P. syringae* RecBCD enzyme recognizes a specific DNA sequence. This specific DNA sequence could potentially be a Chi (*χ*) like sequence; which alters the nuclease activity of RecBCD^Ps^ complex, allowing 3′-ended ssDNA to escape from DNA degradation as observed earlier (2, 8, 23, 24). To further confirm that these DNA fragments are indeed single-stranded, we performed degradation assays in the presence of Exonuclease-I (ExoI), which specifically degrades ssDNA by its 3′→5′ exonuclease activity (25). The RecBCD^Ps^-generated short DNA fragments disappeared in the presence of ExoI (Fig S4A) suggesting that they are indeed ssDNA, similar to Chi-specific DNA fragments produced by *E. coli* RecBCD enzyme after Chi-recognition (8, 23, 24, 26).

Using genetic experiments, we have previously shown that RecBCD^Ps^ enzyme does not recognize *E. coli* Chi sequence (5′-GCTGGTGG-3′) (14, 16). We performed biochemical experiments using pBR322^*χ*+3F3H^ (a pBR322 derivative, containing three tandem *E. coli χ* sequences) (27) and pBR322 (lacking *E. coli χ* sequence) as a DNA substrate. Interestingly, ssDNA fragments produced by RecBCD^Ps^ enzyme were identical in size with both the substrates, confirming that RecBCD^Ps^ does not recognize *E. coli* Chi sequence but recognizes an unknown DNA sequence of pBR322 plasmid (Fig S4B). To locate putative Chi sequence of RecBCD^Ps^ enzyme, we amplified 3.6 kb segment of pBR322 using primer OROPI and OROPII (Supplementary Table ST2), which excludes rop region of the plasmid (Fig. S5A). Amplified fragments were 5′-labeled, and the assays were performed in the presence of excess magnesium. Apparently, all the three shorter ssDNA fragments were also observed with the 3.6 kb DNA substrate, indicating the presence of putative Chi sequence within 3.6 kb of the plasmid (Supplementary Fig. S5C).

### Chi dependent protection of pBR322 DNA fragments are strand specific

*E. coli* RecBCD enzyme recognizes Chi sequence in a specific orientation, 5′-GCTGGTGG-3′, when enzyme enters through 3′-end (28) and the 3′-ended ssDNA is being protected from RecBCD nuclease activity after Chi recognition (8, 29, 30). Here, we examined which strand of the linearized pBR322 is being protected to produce these discrete DNA bands (Fig. 3B). Hence, we performed DNA degradation assay with RecBCD^Ps^, in which only one DNA strand was labeled at a time. We individually labeled the OROPI and OROPII primers with *γ*^32^P at 5′-end and amplified the 3.6 kbp region of pBR322 plasmid with one unlabeled and another ^32^P-labeled primer or with both labeled primers. In our assay, we always used sub-saturating concentration of RecBCD^Ps^ enzyme compared to DNA substrate, so that RecBCD^Ps^ enzyme enters through either one or the other end, but not through both the ends. DNA degradation assays using these DNA substrates produced all the three bands when top strand (OROPII end) was alone labeled, or when the both strands were end-labeled (Fig. 6A). We did not observe any ssDNA band when the bottom strand (OROPI end) was labeled (Fig. 6A), suggesting that RecBCD^Ps^ recognizes a Chi-like sequence in a specific orientation and has a polarity for Chi sequence recognition. These results also suggest that all the Chi-like sequences are on the top strand.

**FIGURE 6.**
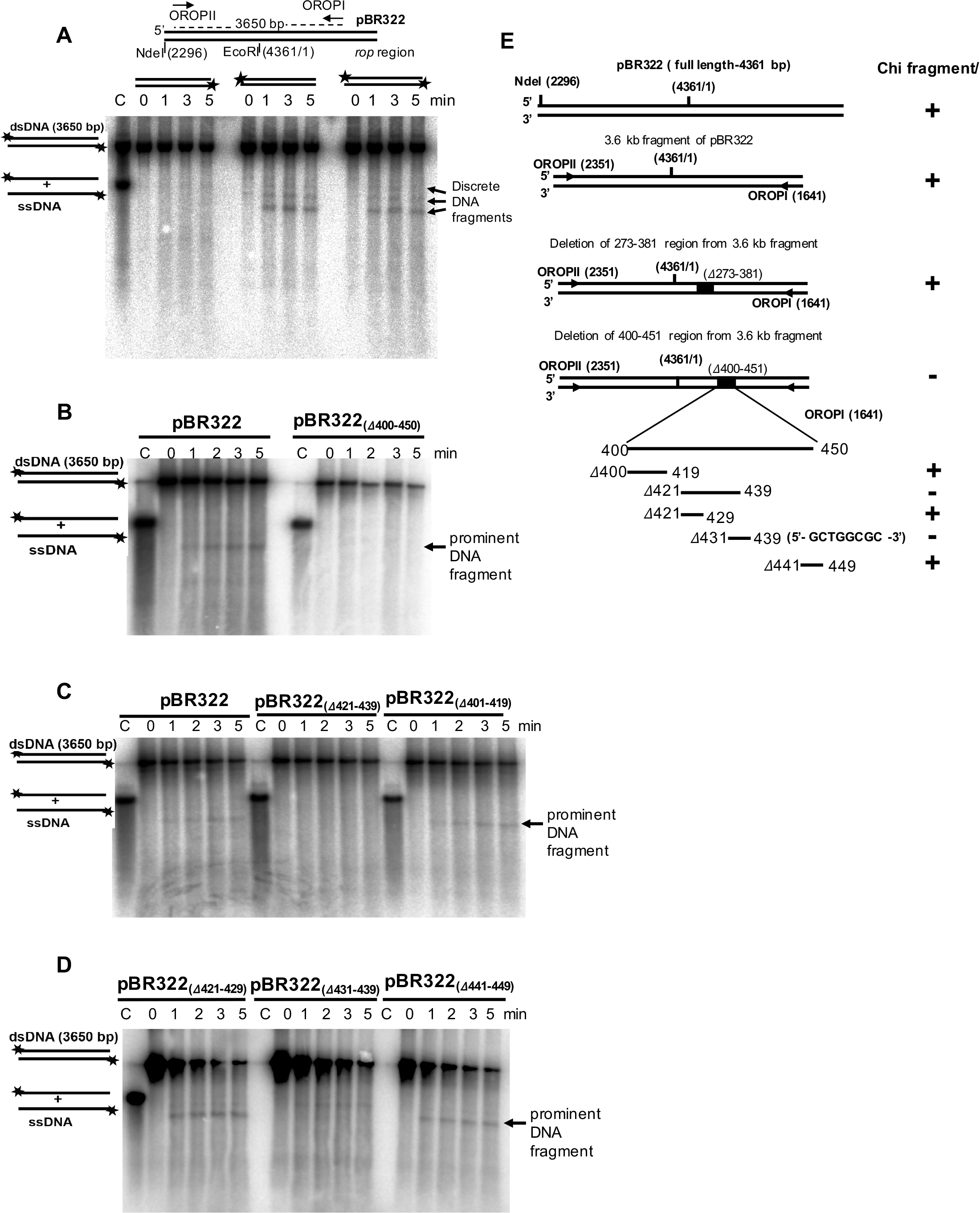
Identification of Chi sequence using PCR amplified pBR322 fragment (3.6 kb) containing internal deletions as a DNA substrate. **(A)** Top panel; Schematic representation of NdeI linearized pBR322 plasmid DNA. The locations of OROPI and OPROPII primers used in PCR amplification of 3.6 kb fragments are indicated. Bottom panel; DNA degradation reactions performed using the PCR amplified 3.6 kb DNA as substrate, having either bottom strand labeled (left panel) or top strand labeled (middle panel) or both strand labeled (right panel). The lane C contains [5′-^32^P] labeled heat-denatured ssDNA as a control. Notably, the top strand labeling of DNA substrate resulted in appearance of discrete DNA fragments. **(B)** Deletion of 400-450 bp of pBR322 (pBR322_(Δ400-450)_) resulted in disappearance of DNA fragments. But, it is clearly visible when intact 3.6 kb pBR322 was used as a substrate, suggesting the presence of putative Chi^Ps^ sequence in this region. **(C)** Further deletion of 401-419; 421-439 bo regions; and, **(D)** 421-429; 431-439; 441-449 bp regions of pBR322 shows that the DNA fragments are indeed from the 431-439 bp region of the pBR322 DNA sequence. **(E)** A schematic representation of 3.6 kb region of pBR322 and deleted regions within, are shown in the left panel. Right panel shows the presence (+) or absence (-) of intense protected band (Chi-like fragments), when these constructs are used as assay substrates.

We then performed DNA degradation assays using two other DNA substrates amplified from the pBR322 plasmid. A PCR amplified 2.8 kbp DNA fragment (using OROPII and pBRS1R primers) and a 2.5 kbp PCR amplified fragment (using OROPII and pBRB1R primers set (Fig. S6A). The DNA band of the lowest size (~2.4 kb) and the highest intensity (the prominent DNA band) was the common ssDNA fragment produced by RecBCD^Ps^ in all the three DNA substrates (Supplementary Fig.S6B). This suggests that one of the Chi-like sequences has the strongest RecBCD-inhibitory activity and this site (we designate it as Chi^Ps^) is located proximal to the 2.5 kbp substrate end (as the protected ssDNA was about ~ 2.4 kb). Two upper DNA bands with less intensity could be due to variants of Chi^Ps^, which might have a weak inhibitory effect on RecBCD^Ps^ nuclease activity (see below).

### Identification of Chi^Ps^ as an 8-mer (5′-GCTGGCGC-3′) sequence that modulates *P. syringae* RecBCD nuclease activity and protects DNA from further degradation

To identify the precise location of Chi^Ps^ site in pBR322 DNA substrate, we generated several internally deleted constructs of pBR322 plasmid by site-directed segment-deletion as described in Materials and methods. The 3.6 kbp DNA region of pBR322 plasmid with deleted region/s were PCR amplified using OROPI and OROPII primers. DNA degradation assays were then performed with RecBCD^Ps^ enzyme using the DNA fragments with internal deletion as substrates (Supplementary Fig. S7).

The localization of Chi^Ps^ on the plasmid was first based on two hypotheses: the sequence might be GC rich and should be located close to the right end of 2.5 kbp DNA fragment. From the pBR322 nucleotide sequence analysis, it appeared that a GC-rich region spanning the region between 273-nucleotides within *tet^R^* gene of pBR322 might contain the Chi^Ps^. Accordingly, 3.6 kbp DNA containing two deletions (Δ273-381 and Δ400-450) were initially tested by the Chi-protection assays.

The 3.6 kbp DNA deleted for 273-381 region produced the Chi-specific high-intensity fragment (Supplementary Fig. S7), while 400-450 nucleotides deletion failed to produce it (Fig. 6B). This indicated that the Chi^Ps^ sequence is located within 400-450 nucleotide region of the pBR322 plasmid. The three additional deletions within the 400-450 region of the plasmid segment were made; Δ401-419, Δ421-439, and Δ441-449 (Fig. 6C, D). DNA degradation assays with the fragments in which these sequences were deleted revealed that Chi is located within the 421-439 nucleotides segment, as these deletions did not produce a protected prominent DNA fragment (Fig. 6C). we further made two more deletion constructs (Δ421-429 and Δ431-439) within the 421-439 nucleotides region, and tested for Chi-specific fragment production. Interestingly, deletion of base-pair from 431-439 bp abolished the prominent DNA band (Fig. 6D) indicating that the deleted region between 431-439 bp, 5′-GCTGGCGCC-3′ contains the putative Chi sequence of *P. syringae.* The schematic representation of all the constructs with deleted nucleotide regions and, the presence or absence of protected prominent DNA band (putative Chi sequence) are also shown in Fig. 6E.

The identification of putative Chi^Ps^ sequence further prompted us to investigate the reason for producing an apparent three distinct DNA fragments by RecBCD^Ps^ enzyme (Fig. 3B). We speculated that pBR322 DNA substrate might have sequences that are similar but not identical to Chi^Ps^ sequence, and might act like Chi^Ps^ sequence with weaker recognition property under the *in vitro* assay conditions. The 9-mer (5′-GCTGGCGCC-3′) sequence appears only at one place on the pBR322 substrate. Considering the *E. coli* Chi sequence is an octamer, we first looked in to 5′-GCTGGCGC-3′ (first 8 nucleotides of the 9-mer from the 5′-end), and 5′-CTGGCGCC-3′ (eliminating first G of the 5′-end) as possible Chi-like sequences. These two octamer sequences appear at a single location on the pBR322 substrate. However, among the 7-mer combinations we looked into, the first 7 nucleotide sequence, 5′-GCTGGGC-3′, was located in three reasonable positions on pBR322 substrate which could potentially produce discrete DNA bands as observed in DNA degradation assays. Hence, we further focused on 5′-GCTGGCGC-3′ sequence as a putative Chi^Ps^ sequence. We found that the putative Chi^Ps^ sequence (5′-GCTGGCGC-3′) is located at 431-438 bp position of pBR322 and, the similar 7-mer sequences at three different locations. The first one at 964-971 position (5′-GCTGGCG**T**-3′); the second one at 1493-1500 position (5′-GCTGGCG**G**-3′) and the third one at 2525-2532 position (5′-GCTGGCG**T**-3′) (Fig S8). All the three-bands show 7 bases identical to the 8 bp Chi^Ps^ sequence (5′-GCTGGCGC-3′), and occur in the same orientation (5′→3′) as the Chi^Ps^. The RecBCD^Ps^ enzyme might recognize 8 mer sequences (5′-GCTGGCGC-3′) as well as the 7+1 mer sequences (5′-GCTGGCG+**T/G**-3′) resulting in multiple DNA bands. Although it is expected to observe four discrete DNA bands, we observed only three DNA bands. The fourth-expected DNA band with a size of ~230 bp (when RecBCD^Ps^ enzyme enters from 3’-side of the NdeI linearized pBR322 substrate as depicted in Fig. S8) was not apparent in in-vitro experiments performed on agarose gels. Possibly, it is the last Chi-like sequence to recognize by RecBCD enzyme and also being the shortest one with ~230 bp (Fig. S8).

To better define the Chi^Ps^ sequence, we mutated the **T** to **C** at the 971^st^ position of pBR322, which creates a sequence identical to the putative 8-mer Chi^Ps^ sequence and performed the Chi-protection assay with RecBCD^Ps^ enzyme. Interestingly, the intensity of the Chi-protected second DNA fragment (after T to C mutation) appeared stronger, and the intensity was similar to the prominent DNA band (Fig. 7A), which establish the strong recognition of octamer sequence (5′-GCTGGCGC-3′) as Chi^Ps^.

**FIGURE 7.**
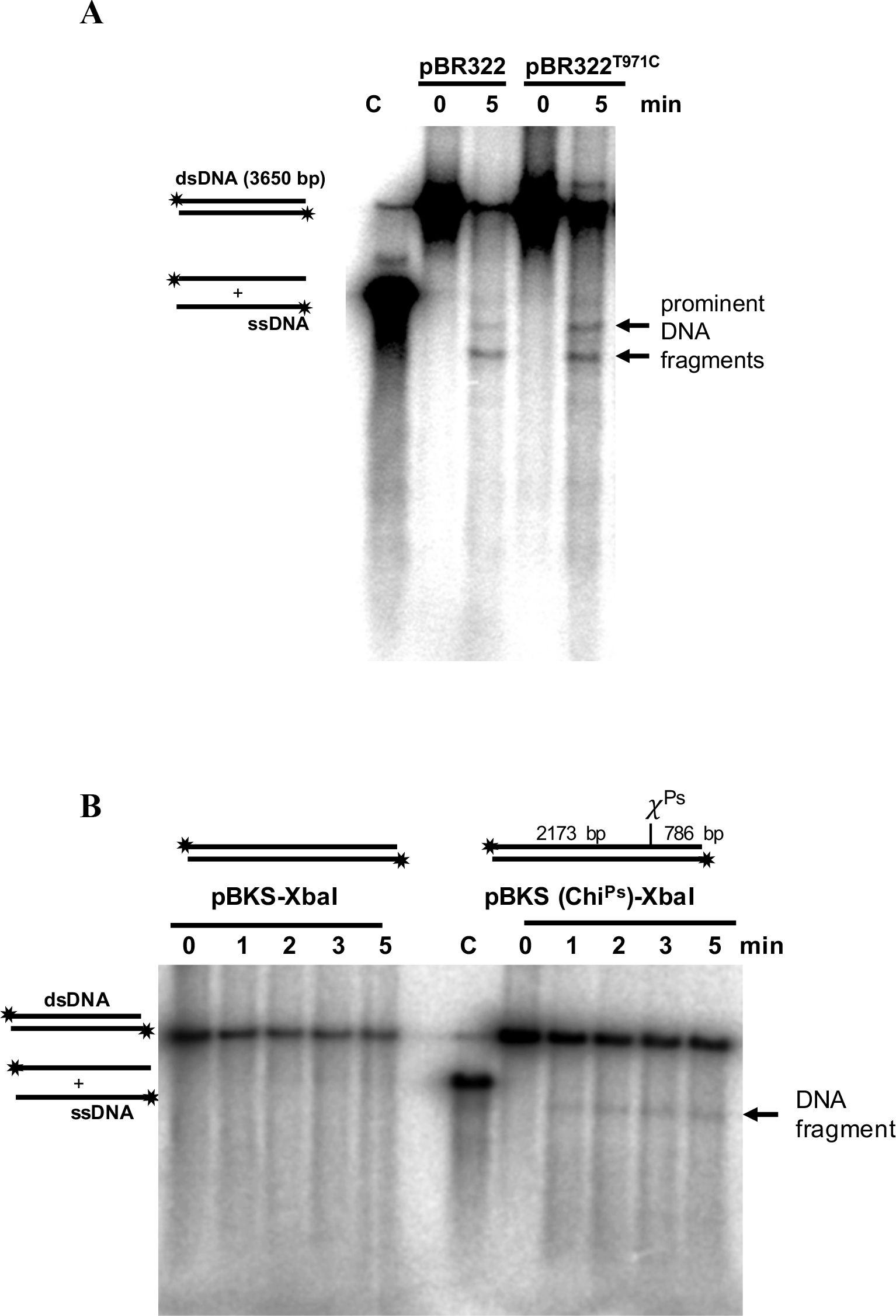
(A) DNA degradation of modified fragment of pBR322 by RecBCD^Ps^ enzyme. Apart from one 8-mer Chi^Ps^ sequence at 431th position of pBR322, there are two 7-mer similar sequences present in the plasmid pBR322. One such similar sequence at 964^th^ position (5′ GCTGGCG**<u>T</u>** 3′) was mutated to make it an 8-mer sequence (5′ GCTGGCG**<u>C</u>** 3′). This substrate (pBR322^T971C^) that has two Chi sequences in the same orientation when used as a substrate, yielded two DNA fragments with high intensity. **(B) The DNA degradation pattern of** *XbaI* **digested linear dsDNA of pBKS plasmid by RecBCD^Ps^ enzyme.** Chi^Ps^ are inserted into pBluescript vector (pBKS) by site directed insertion. *XbaI* linearized pBKS vector or pBKS with Chi^Ps^ was 5′-end labeled with ^32^P has been used for assays. The discrete ssDNA of expected size was produced when pBKS (Chi^Ps^) was used as a substrate, whereas it is absent, when native pBKS was used.

### Cloning of Chi^Ps^ in pBKS plasmid and confirmation of ectopic Chi^Ps^ activity

To further confirm that the octamer sequence (5′-GCTGGCGC-3′) is indeed the Chi sequence in *P. syringae* and can function as an ectopic Chi^Ps^ sequence, we cloned the octanucleotide sequence into pBKS which is devoid of Chi^Ps^ sequence (Table ST1). RecBCD^Ps^ reactions under excess magnesium were performed with XbaI digested linear pBKS (Chi^0^) plasmid and the Chi^Ps^ containing pBKS (pBKS-Chi^Ps^)plasmid. As expected, no Chi-protected DNA bands were observed using the linear pBKS (Chi^0^) DNA as substrate. In contrast, the expected Chi specific DNA fragment was observed in the case of XbaI digested pBKS-Chi plasmid (Fig. 7B). These experiments confirm that RecBCD^Ps^ recognizes an 8-mer sequence (5′-GCTGGCGC-3′) as Chi^Ps^ sequence during resection of double-stranded DNA ends, and thus Chi^Ps^ attenuates the nuclease activity of RecBCD enzyme and promotes DNA recombination and repair.

## DISCUSSION

We have earlier shown that RecBCD protein complex is essential for cold adaptation in Antarctic *P. syringae* Lz4W (14-16). In this study, we have analyzed the biochemical properties of the wild-type (RecBCD^Ps^) and three mutant enzymes (RecB^K28Q^CD, RecBCD^K229Q^, and RecB^D1118A^CD) to analyze the role of RecBCD during growth at low temperature. We also report for the first time, *Pseudomonas* octameric Chi-sequence (Chi^Ps^), (5′-GCTGGCGC-3′), and its ability to attenuate the nuclease activity of *P. syringae* RecBCD enzyme *in vitro.*

### Role of ATPase activity of RecB and RecD subunits

The analyses of wild-type and mutant enzymes of *P. syringae* have revealed that mutation in the critical ATP binding sites of RecB (RecB^K28Q^) and RecD (RecD^K229Q^) motor subunit reduces the ATPase activity of trimeric RecBCD complex by ~10 fold. The mutation in the nuclease active-site does not affect the ATPase activity. *P. syringae* cells carrying the alleles of RecB^K28Q^ or RecD^K229Q^ are sensitive to cold temperature, UV irradiation and MMC (14). The ~ 10-fold reduction of ATPase activity observed *in-vitro* supports the idea that ATP hydrolysis by RecB and RecD subunits in RecBCD^Ps^ holoenzyme is essential for *P. syringae* survival at low temperature.

### DNA unwinding and degradation activities of RecBCD and mutant enzymes

Our data suggest that the DNA unwinding and degradation properties of RecBCD enzyme depend on ATP and Mg^++^ concentrations, and on the temperature *in vitro.* Under limiting magnesium reaction condition, at 22°C, the RecB^K28Q^CD, RecBCD^K229Q^ and RecB^D1118A^CD enzymes have retained about 4%, 77% and 85% of the wild-type DNA unwinding activity respectively, while at 4°C, we could not detect the DNA unwinding activity of both wild-type and mutant RecBCD^Ps^ enzymes under the limiting magnesium reaction condition. In contrast, when magnesium was in excess over ATP, the DNA unwinding and degradation activity of the RecBCD enzymes could be measured 22°C and 4°C). These results suggest that the excess of magnesium over ATP is favorable for the helicase and nuclease activities of RecBCD^Ps^ enzyme.

The RecB^K28Q^CD enzyme is a poor helicase and with no detectable nuclease activity *in vitro* even at 22°C. However, at 4°C, RecB^K28Q^CD enzyme failed to unwind and degrade the DNA under both limiting and excess magnesium reaction conditions. Hence, low-temperature growth sensitivity of the *P. syringae recB^K28Q^CD* mutant can be attributed to the lack of RecBCD unwinding and/degradation activities at 4°C.

The RecBCD^K229Q^ enzyme with defective RecD ATPase unwinds and degrades the dsDNA, albeit at the reduced rate. The DNA unwinding/degradation rate of RecBCD^K229Q^ mutant enzyme is reduced to one-third of the wild-type enzyme at 4°C. Therefore, we propose that its inability to support the DNA repair process and growth at low temperature is possibly due to its decreased DNA unwinding/degradation activity, particularly at low temperature. Interestingly, the RecBCD^K229Q^ enzyme shows lack of discrete DNA fragments (putative Chi-specific fragments) production. However, the *E. coli* RecD ATPase defective RecBCD enzyme produces the Chi-specific fragments *in vitro* and confers DNA repair proficiency *in vivo* (31). This converse observation suggests that the RecD subunit of RecBCD^Ps^ enzyme has a distinct role in DNA repair and recombination function in *P. syringae.* Despite both RecBCD^K229Q^ and RecB^K28Q^CD mutant enzymes being similarly defective in ATP hydrolyzing activity, their DNA unwinding activity was largely varied. This possibly be due to the selective motor dependency of the enzyme complex, similar to observation made in *E. coli* RecBCD enzyme (31), in which, the RecB motor is the absolute requirement for Chi recognition and the motor activity of RecBCD complex.

The RecB^D1118A^CD enzyme is a processive helicase with no detectable nuclease activity *in vitro,* at both 22°C and 4°C. Interestingly, *P. syringae* cells carrying *recB^D1118A^* allele are capable of growing at low temperature, suggesting nuclease activity is not essential for its growth at low temperature (14).

### RecF pathway role in RecBCD nuclease defective strain and its importance in *P. syringae* growth at low temperature

In *E. coli,* the RecF pathway along with RecJ (5′→3′ ssDNA specific nuclease) is known to work with nuclease-defective, RecA loading-deficient RecBCD (32). The role of RecBCD nuclease defective - RecF hybrid pathway in *P. syringae* is also evidenced. Deletion of *recF* gene alone (in WT cells) is cold resistant (15). However, deletion of the *recF* gene in *recB^D1118A^* mutant renders cold sensitivity (unpublished observation, Apuratha T Pandiyan and Malay K Ray). The over-expression of RecJ in pRecB^Δnuc^CD (RecB nuclease domain deleted) expressing *P. syringae recBCD* null strain alleviated the slow growth phenotype of *P. syringae* cells at low temperature (14), suggesting RecJ role in nuclease-defective RecBCD cells. These observations indicate RecFOR-RecJ role in RecB nuclease-defective *P. syringae* strain.

*recA* deleted *P. syringae* cells grow slowly at low temperature (15). This suggests that RecBCD enzyme alone, in the absence of RecA, can rescue low temperature-induced replication fork arrest possibly by suppressing chromosomal lesions via DNA degradation of reversed replication fork (33). Interestingly, the combination of a *recA* deletion and a nuclease defective RecBCD mutation (*recB^D1118A^CD*) causes cold sensitivity (15), suggesting a direct role of RecA in rescuing low temperature induced replication fork arrest in a RecB nuclease defective strain. Therefore, we propose that, in RecB nuclease defective strain, the RecF pathway enables RecA mediated DNA repair and thus, protects cells from low temperature induced DNA damage.

### Identification and characterization of Novel octameric Chi^Ps^ sequence

One of the novel findings in this study is the identification of *P. syringae* Lz4W Chi-sequence (Chi^Ps^), 5′-GCTGGCGC-3′. This sequence has not been identified so far in any *Pseudomonas* species. Most of the identified bacterial Chi (*χ*) sequences are GC rich sequences (Table 3) and the number of nucleotides in *χ* sites vary from 5-mer (in *Bacillus subtilis*) to 8-mer (in *E. coli, L. lactis* and *H. influenza*) (34). A study using cell lysate from Pseudomonads indicated that *Pseudomonas* species do not recognize *E. coli χ* sequence (24). We have confirmed this observation earlier by genetic experiments (14), and in the present study, we have biochemically identified the Chi^Ps^ (5′-GCTGGCGC-3′) sequence; which is identical up to bases from the 5′-end to the *E. coli* Chi (Chi^Ec^) (5′-GCTGGTGG-3′) sequence. 7-mer (5′-GCTGGCG-3′) sequence, with change in the last 8^th^ nucleotide, are partially recognized. The mutation of 7-mer sequence to make a perfect 8-mer Chi^Ps^ sequence enables it to be strongly recognized by RecBCD^Ps^. Interestingly, appearance of three ssDNA fragments indicates that RecBCD^Ps^ enzyme can sometimes bypass Chi^Ps^ and recognize the next putative Chi-like sequence. Similar observations were made in *E. coli,* where the probability of Chi recognition by *E. coli* RecBCD enzyme and nicking the DNA is about 30-40% (28, 35).

**TABLE 3.** Chi sequences identified in different bacteria.

| Bacteria | Chi sequence | Recombination machinery |
| --- | --- | --- |
| *Escherichia coli* | 5' GCTGGTGG 3' | RecBCD (43) |
| *Bacillus subtilis* | 5' AGCGG 3' | AddAB (44) |
| *Lactococcus lactis* | 5' GCGCGTG 3' | RexAB (34) |
| *Streptococcus pneumoniae* | 5' GCGCGTG 3' | RexAB (45) |
| *Staphylococcus aureus* | 5' GAAGCGG 3' | - (45) |
| *Streptococcus agalactiae* | 5' GCGCGTG 3' | - (45) |
| *Streptococcus thermophilus* | 5' GCGCGTG 3' | - (45) |
| ***Pseudomonas syringae* Lz4W** | **5' GCTGGCGC 3'** | RecBCD (14) |

Interestingly, the difference between *E. coli* and *P. syringae* RecBCD enzymes are confined to the last three nucleotides (5′- GCTGG<u>TGG</u>-3′ vs 5′- GCTGG<u>CGC</u> -3′). The recent study on the molecular determinants responsible for the Chi recognition by RecBCD enzyme has revealed the importance of RecC channel in Chi recognition. Among the 35 amino acid residues of RecC channel examined, the Q38, T40, Q44, L64 W70 D133, L134, D136, D137, R142, R186 and D705 residues of *E. coli* RecC subunit have shown to affect the Chi recognition property of RecBCD^Ec^ enzyme (36). Surprisingly, all these residues are well conserved in the *P. syringae* RecC subunit. Therefore, the RecC amino acids responsible for recognition of the last three nucleotides of are still elusive. Further analysis of *E. coli* and *P. syringae* RecC subunits could shed more insight on the molecular determinants responsible for the recognition of last three nucleotides of Chi sequence in *E. coli.*

*E. coli* contains 1,008 Chi sequences (37). They are four-to eightfold more frequent than expected by chance and appear on average once every 4.5 kbp. 75% of Chi sites are skewed towards the replicative leading strand in *E. coli* (37) keeping with their function in stimulating double-strand break repair upon replication fork collapse. These observations suggest a role for the RecBCD enzyme as a repair factor functioning towards re-establishment of DNA replication fork in case of collapse. However, this skewed nature is not applicable for *B. subtilis, S. aureus* and *H. influenzae,* where the skew is statistically insignificant, and the Chi sequence (of the respective species) is distributed all over the genome (34). The search for Chi^Ps^ sequence (5′- GCTGGCGC-3′) in the draft genome sequence of *P. syringae* Lz4W (4.98 Mb in 42 contigs, **Accession no. <u>AOGS01000000</u>)** revealed that it contains 1541 Chi^Ps^ sequences (5′-GCTGGCGC-3′) and 4564 of 7-mer Chi^Ps^ sequences (5′-GCTGGCG-3′). The Chi^Ps^ sequence appears once in every ~3 kb and is overrepresented compared to other random octamer sequences. No skewed Chi^Ps^ sequence distribution was observed in *P. syringae* Lz4W genome (A. Pandiyan and M. K. Ray, unpublished observation). Analysis of the closely related *P. fluorescens* Pf0-1 genome **(Accession no. <u>NC 007492.2</u>)** (38) revealed that it contains 2241 Chi^Ps^ sequences and the Chi^Ps^ sequence appears once in every 3 kb and, is almost equally distributed on both strands (1119 Chi-Ps vs. 1122 complementary Chi-Ps sequences). Thus the pattern of Chi-distribution is not universal, and although Chi^Ps^ is over-represented in *Pseudomonas* genome, the orientation bias is not observed.

### Biochemical properties of RecBCD enzyme and its role in *P. syringae* growth at low temperature

This study has revealed the biochemical properties of *Pseudomonas* RecBCD enzyme. The biochemical properties of RecBCD^Ps^ enzyme, compared to the RecBCD^Ec^, are particularly associated with the RecD subunit. In *E. coli,* RecD is dispensable for DNA repair process. The *recD* null *E. coli* strain is hyper-recombinogenic (39) and RecBC^Ec^ enzyme (without RecD) unwinds dsDNA and loads RecA constitutively in a Chi-independent manner (30). Also, the *E. coli* RecBCD enzyme with a mutation in the ATP binding site of RecD subunit produces Chi-specific fragments and cells expressing the mutant enzyme are UV resistant (31). In contrast, *P. syringae,* RecD is essential for the RecBCD’s function (14) and ATP hydrolyzing activity of RecD motor is an absolute requirement for Chi^Ps^ fragments production. Also, cells expressing RecD ATPase mutant enzyme are cold sensitive, UV and MMC sensitive (13). Importantly, Chi-like octameric sequence (5′-GCTGGCGC-3′) attenuates nuclease activity of RecBCD^Ps^ enzyme producing Chi^Ps^ containing ssDNA fragments and thus, can act as a Chi sequence for RecBCD enzyme of *Pseudomonas* species.

Based on our results we propose a model (Figure 8) which explains the role of RecBCD and collaborative DNA repair pathways in rescuing the replication fork arrests in *P. syringae* Lz4W at low temperature. In this model, RecBCD enzyme can rescue replication fork from the arrest by linear chromosomal DNA degradation in a recA-independent manner. When nuclease activity is compromised, the nuclease-defective RecBCD enzyme acts with the *recF* pathway to ensure DNA repair by homologous recombination. In this model, we also propose that motor activity of RecBCD enzyme is essential for rescuing *P. syringae* cells from low temperature induced replication fork arrest.

**FIGURE 8.**
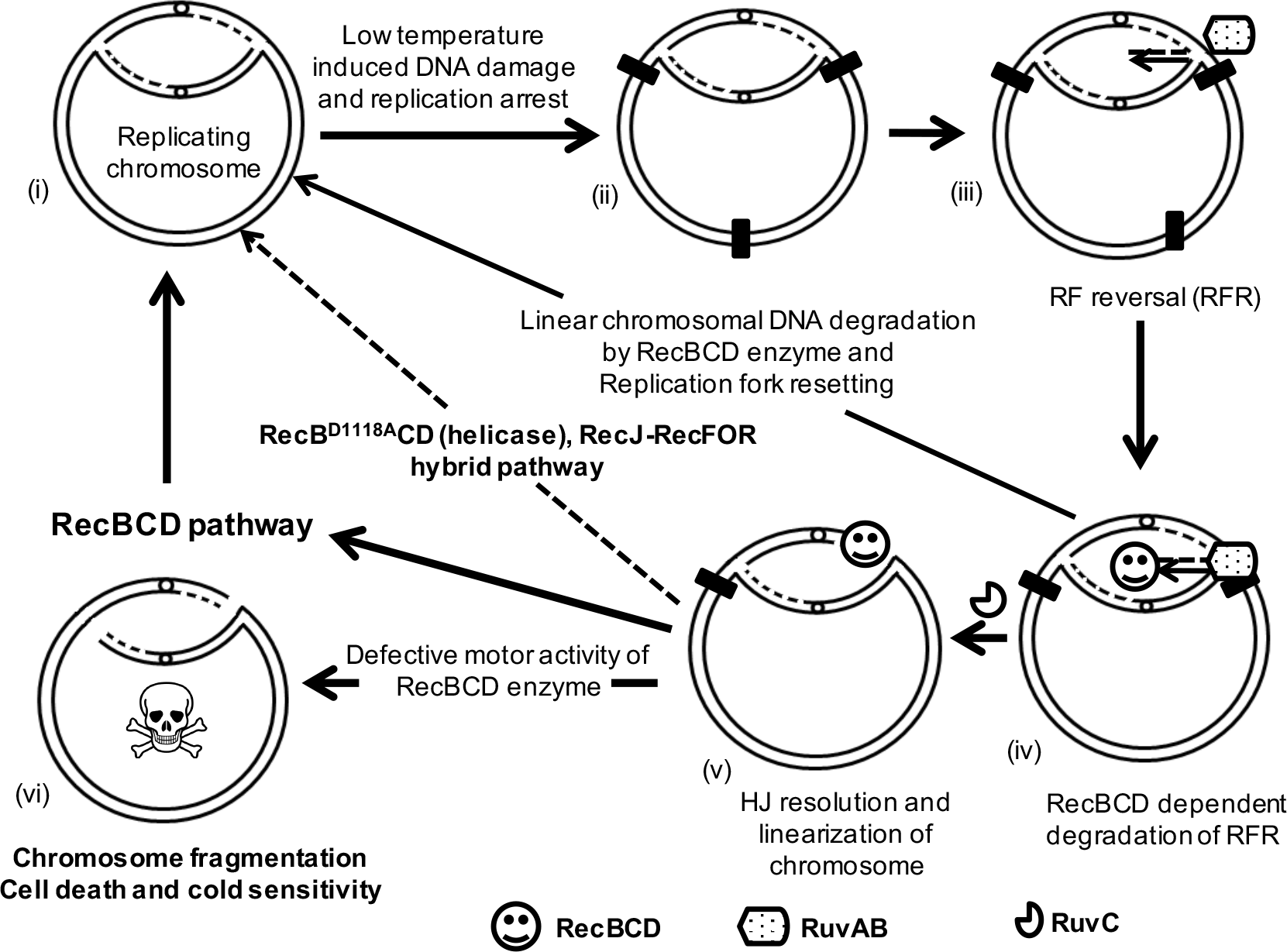
Role of RecBCD dependent DNA repair pathway in rescue of low temperature induced replication forks arrest. (i) A chromosome replicating bi-directionally, (ii) encounters low temperature-induced chromosomal lesion or blockage, (iii), causing replication fork arrest and fork reversal (RFR). RFR is suppressed by linearized chromosomal DNA degradation by RecBCD enzyme and resetting of replication fork. (iv) RFR is stabilized further by RuvAB complex and, (v) further, resolved by RuvC leading to chromosomal linearization. Linearized chromosome is processed by either RecBCD alone or in conjunction (when RecBCD is nuclease defective) with recFOR-recJ hybrid pathway. The defective motors activity of RecBCD enzyme leads to chromosomal fragmentation and cell death at the low temperature.

## MATERIALS AND METHODS

### Bacterial strains, plasmids and growth conditions

The bacterial strains and plasmids used in this study are listed in Tables ST1. The psychrophilic *P. syringae* Lz4W was isolated from a soil sample of Schirmacher Oasis, Antarctica (40) and routinely grown at 22 or 4°C (for high and low temperatures respectively) in Antarctic bacterial medium (ABM) composed of 5 g peptone and 2.0 g yeast extract per liter, as described earlier (16). *E. coli* strains were cultured at 37°C in Luria–Bertani (LB) medium, which contained 10 g tryptone, 5 g yeast extract and 10 g NaCl per liter. For solid media, 15 g bacto-agar (Hi-Media) per liter was added to ABM or LB. When necessary, LB medium was supplemented with ampicillin (100 µg ml^−1^), kanamycin (50 µg ml^−1^), gentamicin (15 µg ml^−1^) or tetracycline (20 µg ml^−1^) for *E. coli.* For *P. syringae,* the ABM was supplemented with tetracycline (20 µg ml^−1^), kanamycin (50 µg ml^−1^) as needed.

pBR322 plasmid DNA (4.3 Kb) and *χ*+-3F3H dsDNA (a pBR322 derivative, containing two sets of three tandem repeats of *χ* sequences (27)) were purified using a Qiagen midi kit. Plasmids were linearized with NdeI restriction endonuclease, dephosphorylated with SAP. The dephosphorylated linear dsDNA was 5′- labeled with T4-PNK and [*γ*-^32^P] ATP as per the manufacturer guidelines. DNA concentrations were determined by absorbance at 260 nm using molar extinction co-efficient of 6500 M^−1^ cm^−1^ (in nucleotides). All restriction enzymes, DNA ligase T4 Polynucleotide Kinase (T4 PNK), Shrimp alkaline phosphatase (SAP) and *E. coli* SSB were purchased from New England Biolabs (MA, USA). Accuprime Pfx DNA polymerase was purchased from Novagen (WI, USA).

### Antibodies and Western analysis

Production of anti-RecB, anti-RecC and anti-RecD antibodies has been described (14). For Western analysis, proteins were separated by SDS-PAGE, transferred onto Hybond C membrane (Amersham Biosciences), and probed with appropriate antibodies. The immunoreactive protein bands were detected by alkaline phosphatase-conjugated anti-rabbit goat antibodies (Bangalore Genie, India). For quantification, the blots were scanned with a HP scanjet and band intensities were measured using Image J software (rsbweb.nih.gov/ij/).

### Over expression and purification of recombinant proteins

The LCBD (Δ*recBCD*) strain harboring pGHCBD, pGHCB^K28Q^D pGHCBD^K229Q^ pGHCB^D1118A^D plasmids (14) were initially grown in 10 ml ABM broth containing kanamycin (50 µg/ml) for 24 hrs at 22°C. 1% of above culture was inoculated into a 2-liter conical flask containing 500 ml ABM broth with kanamycin (50 µg/ml). The culture was then incubated at 22°C with aeration for 24 hrs. Later, the bacterial cells were harvested by centrifugation at 4°C, 6000 rpm for 10 min. The bacterial cell pellets were stored at −80°C. The cells pellet was removed as and when required for the purification of proteins.

All the recombinant proteins in this study were expressed with His-tag on N-terminus of RecC and purified by nickel-nitrilotriacetic acid (Ni-NTA) affinity chromatography as described in the manufacturer’s protocol (Qiagen, New Delhi, India). In brief, cell lysate of overexpressed strains containing His-tagged proteins were prepared by dissolving the cell pellet in 10 ml lysis buffer (50 mM NaH_2_PO_4_ (pH 7.4), 300 mM NaCl, 10 mM imidazole and 10% glycerol) and lysed by mild sonication. The sonicated cell lysate was then centrifuged with 14,000 rpm at 4°C, for 30 min to remove insoluble cellular debris. The supernatant was passed through a pre-equilibrated column containing 1 ml slurry of Ni-NTA agarose beads and column was allowed to bind Ni-NTA agarose beads with His-tagged proteins. Further, column was washed with 4-5 volumes of wash buffer (50 mM NaH_2_PO_4_ (pH 7.4), 300 mM NaCl, 20 mM imidazole, and 10% glycerol). Finally, the bound proteins were eluted with 1-2 ml elution buffer (50 mM NaH_2_PO_4_(pH 7.4), /300 mM NaCl / 300 mM imidazole /10% glycerol).

Gel filtration technique (size-exclusion column chromatography) was employed for the purification of His-tagged RecBCD complex and the mutant protein complexes. For this, Superose-gel filtration column (Pharmacia Fine Chemicals) was used in the fast protein liquid chromatography (FPLC) (Pharmacia Fine Chemicals). Initially, the column was pre-equilibrated with buffer contained 20 mM Tris HCl (pH 7.5), 0.1 mM EDTA, 150 mM NaCl, 0.1 mM PMSF and 10% glycerol. Later, 0.5 ml of Ni-NTA purified protein solution was injected to the column and allowed to pass through the column at a flow rate of 0.4 ml/min. Optical density at 280 nm was recorded for the eluted protein fractions and the fractions were collected in separate microcentifuge tubes. The protein fractions were then analyzed on SDS-PAGE stained with coomassie brilliant blue or silver nitrate. The gel filtration protein fractions of interest were further membrane dialyzed in 50% glycerol containing gel filtration buffer. RecBCD enzyme concentrations were determined by measuring OD_280_ and using molar extinction co-efficient 4.7 × 10^5^ M^−1^ cm^−1^ as determined by ExPASy – ProtParam tool (https://web.expasy.org/protparam).

### Thin-layer chromatography based assay

ATPase activity of RecBCD and mutant proteins was assayed at different temperatures by thin layer chromatography (TLC) on polyethylene-imine (PEI)-cellulose plates (E-Merck, Germany). The assay was performed as described earlier (41, 42). Assays were carried out at indicated temperatures in a reaction volume of 20 µl containing 25 mM Tris acetate (pH 7.5), 1 mM Mg acetate, 1 mM DTT, 100mM NdeI linearized pBR322-dsDNA and 200 µM ATP as a substrate with 2 nM RecBCD and mutant enzymes. One µl of 100 times diluted 10 mCi ml^−1^ stock of [*γ*-^32^P] ATP (specific activity 3000 Ci mmol^−1^) was used as a tracer in each reaction to measure the rate of ATP hydrolysis. Following 0, 1, 2, 3, 5 10 minutes of reaction, 0.5 µl aliquots of the samples were spotted on TLC plate, air-dried and were allowed to develop in a mobile phase containing 0.5 M formic acid and 0.5 M lithium chloride for 15 minutes. The TLC plate was dried and exposed to the Phosphor imaging plate for 4-6 hrs. The Imaging plates were scanned in a Phosphor Imager, and the amounts of ^32^Pi and [*γ*-^32^P] ATP were quantified using Image gauge software (Fuji-3000). Further, data were analyzed using GraphPad Prism 4.0 software.

### DNA unwinding assay

Plasmid DNAs were linearized with appropriate restriction enzymes in the presence of shrimp alkaline phosphatase and 5′-end-labeled by T4 polynucleotide kinase and [*γ*-^32^P] ATP. Subsequent purification of labeled DNA was accomplished by passage through a MicroSpin S-200 HR column (Amersham biosciences-GE healthcare, Buckinghamshire, UK). The reaction mixtures contained 25 mM Tris acetate (pH 7.5), 2 mM magnesium acetate (as indicated), 1 mM DTT, 10 µM (nucleotides) linear pBR322 dsDNA P^32^-labeled at 5′- end, 5 mM ATP, 2 µM *E. coli* SSB protein and 0.5 nM RecBCD^Ps^ or mutant enzymes. DNA unwinding reactions were started with the addition of either enzyme or ATP, after pre-incubation of all other components at 22 or 4°C for 5 min. Assays were stopped at the indicated times by addition of proteinase K to a final concentration of 0.5 mg/ml, which was dissolved in sample loading buffer (125 mM EDTA, 40% glycerol, 2.5% SDS, 0.25% bromophenol blue, and 0.25% xylene cyanol). After 5-min incubation with proteinase K at room temperature, the reaction products were run on a 1% (w/v) agarose gel in a 1X TBE (45 mM Tris borate (pH 8.3) and 1 mM EDTA) buffer at 25-30 constant volts for 15 hrs. Agarose gels were dried, exposed to phosphor imaging plates and quantified using Phosphor Imager (Fuji-3000). Further, data were analyzed using image gauge software.

The DNA unwinding rates of were measured by using the following formula

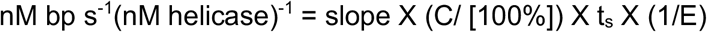

Where C is the concentration of linear dsDNA substrate in base-pairs in nM (i.e., 5000 nM), t_s_ is the time in seconds, and E is the enzyme concentration in nM.

### DNA degradation assay

The assays were performed as described above in DNA unwinding assay, except that the reaction mixtures contained 6 mM magnesium acetate and 2 mM ATP.

### Single-stranded DNA endonuclease assay

The endonuclease activity of RecBCD enzyme on ssDNA was examined using a circular M13 ssDNA substrate as described previously (20, 22). In brief, endonuclease activity was tested in 3 different buffer conditions. The first reaction mixture contained 50 mM MOPS (pH=7.5), 1 mM ATP, 10 mM MgCl_2_, 4.16 nM circular M13 ssDNA with 0.5 nM RecBCD. The second reaction mixture contained 25 mM Tris-acetate, 1 mM ATP, 8 mM Mg-acetate, 1 mM DTT, 4.16 nM M13 ssDNA with 0.5 nM RecBCD. The third reaction mixture contained 25 mM Tris-acetate, 2 mM ATP, 6 mM Mg-acetate, 1 mM DTT, 4.16 nM M13 ssDNA with 0.5 nM RecBCD. After the reaction, samples were removed at the indicated times; quenched with 120 mM EDTA, 40% (v/v) glycerol, and 0.125% bromphenol blue; and loaded on a 0.8% agarose gel in 1X TBE (90 mM Tris borate, 2 mM EDTA). The gel was run at 4 V/cm for 3 h and stained with ethidium bromide (0.5 mg/ml). The bands were visualized by exposure to UV light.

### Site directed deletion of regions from plasmid pBR322

Different internal regions of plasmid pBR322 were deleted using site directed deletion. In short, primers were designed consists of sequences flanking the region to be deleted. PCR was performed to amplify whole Plasmid DNA using Accuprime Pfx DNA polymerase (Invitrogen). After PCR, reaction mixture was subjected for the overnight DpnI digestion. 5 µl of reaction mixture was then transformed into DH5a ultra-competent cells. All selected regions, which had to be deleted was in tetracycline resistance gene of the plasmid. Therefore, for primary screening only those colonies were selected which were Amp^R^ and Tet^s^. Deletion was further confirmed by PCR and sequencing. All primer list and corresponding deleted regions are shown in Table ST2.

## ACCESSION NUMBERS

**Accession no. AOGS01000000,** the draft genome sequence of *P. syringae* Lz4W and,

**Accession no. NC_007492.2,** the genome sequence *Pseudomonas fluorescens* Pf0-1.

## SUPPLIMENTARY MATERIALS

Supplementary material related to this manuscript is attached.

## ACKNOWLEDGEMENTS

Research in M.K.R. laboratory is supported by the Council of Scientific and Industrial Research (CSIR), Government of India. A part of the work was supported by a grant to M.K.R. from Department of Science and Technology (DST), Government of India. T.L.P and A.K.S. acknowledge CSIR, India for research fellowships. We also thank Prof. Stephen C Kowalczykowski, Prof. Benedicte Michel Benedicte and Dr. Naofumi Handa for their critical reading and comments on the manuscript.

## AUTHOR CONTRIBUTIONS

Theetha L. Pavankumar & Anurag Kumar Sinha both have contributed equally to this work.

T.L.P., A.K.S. and M.K.R designed the experiments. T.L.P. and A.K.S. conducted the experiment.

T.L.P., A.K.S. and M.K.R analyzed the results and wrote the manuscript.

